## Supplemental Figures for "Palmitoylated Importin α Regulates Mitotic Spindle Orientation Through Interaction with NuMA"

**Expanded View Figures**

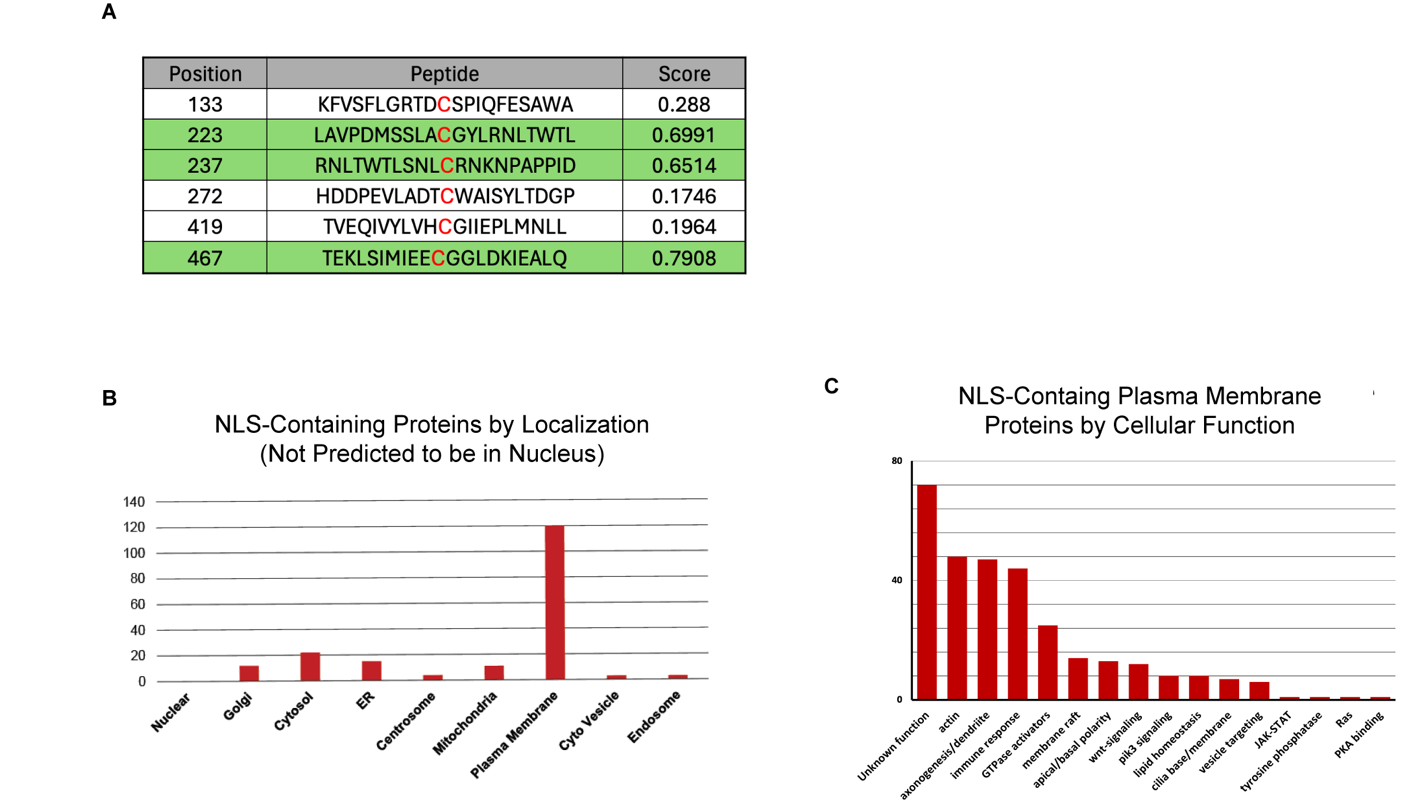

**Expanded View Figure 1. Proteome Screens Confirm Palmitoylation of Human Importin α-1 and Enrichment of NLS Containing Proteins by Cellular Localization and Function.**

A) Prediction of palmitoylated cysteine residues in Human Importin α-1 (KPNA2) by GPS-Palm. Prediction score is on a 0-1 scale with 0 representing low confidence of palmitoylation and 1 representing high confidence of palmitoylation. Three highest confidence residues are highlighted in green.

B) Cellular localization of NLS containing proteins not predicted to be in the nucleus.

C) Cellular functions of NLS containing proteins found at the PM sorted by gene ontology terms.

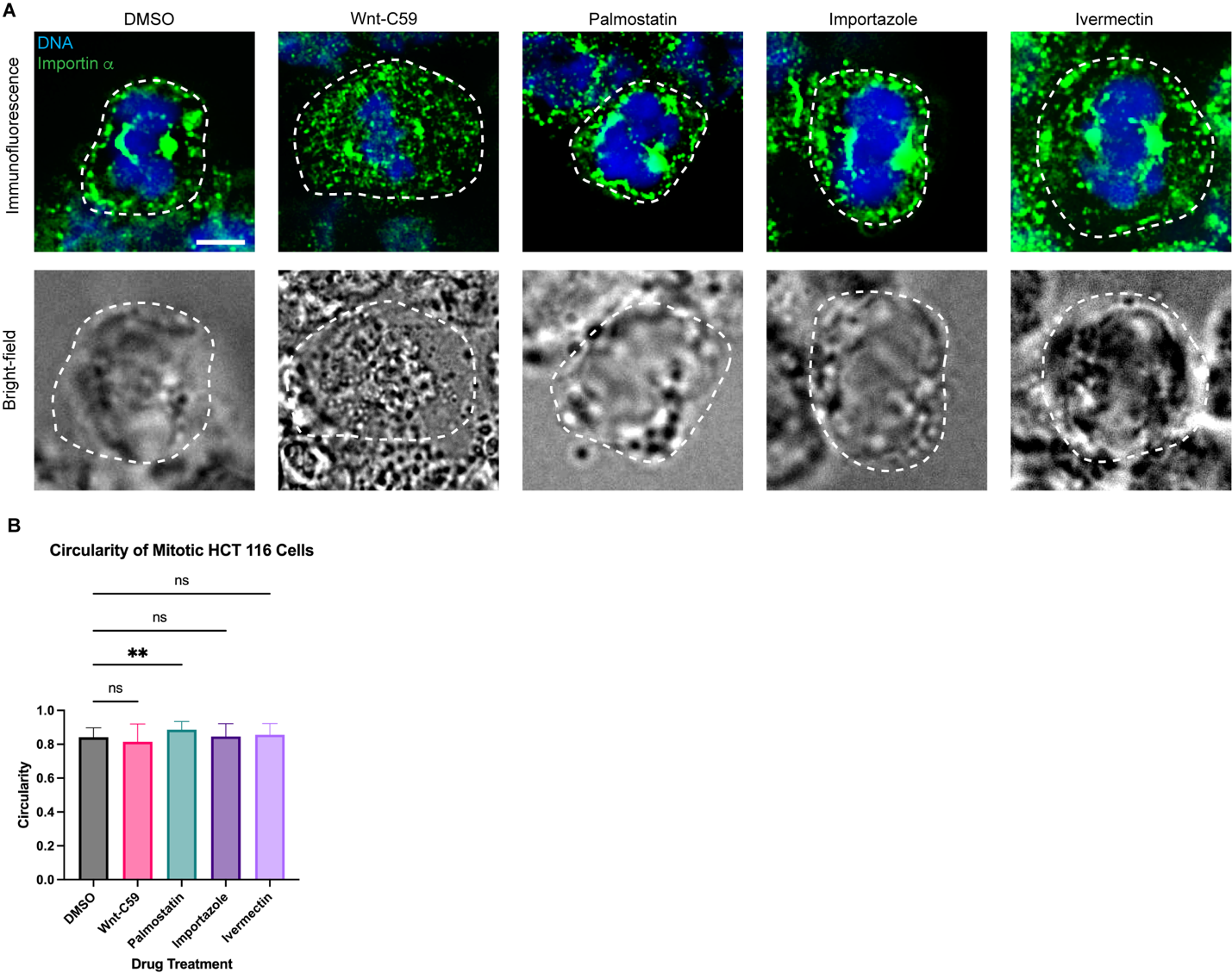

**Expanded View Figure 2. KPNA2 Cellular Localization Boundary Determination.**

A) Representative immunofluorescence and bright-field images for cells used in Figure 1A to determine KPNA2 localization and cell boundary determination for each drug treatment. Scale bar=5μm.

B) Quantification of the circularity of metaphase-arrested HCT116 cells treated with DMSO, 10μM Wnt-C59, 50μM palmostatin, 40μM importazole or 25μM ivermectin for 1 hour. Mean +/- SEM n=60, **p<0.01 determined by Student’s t-test.

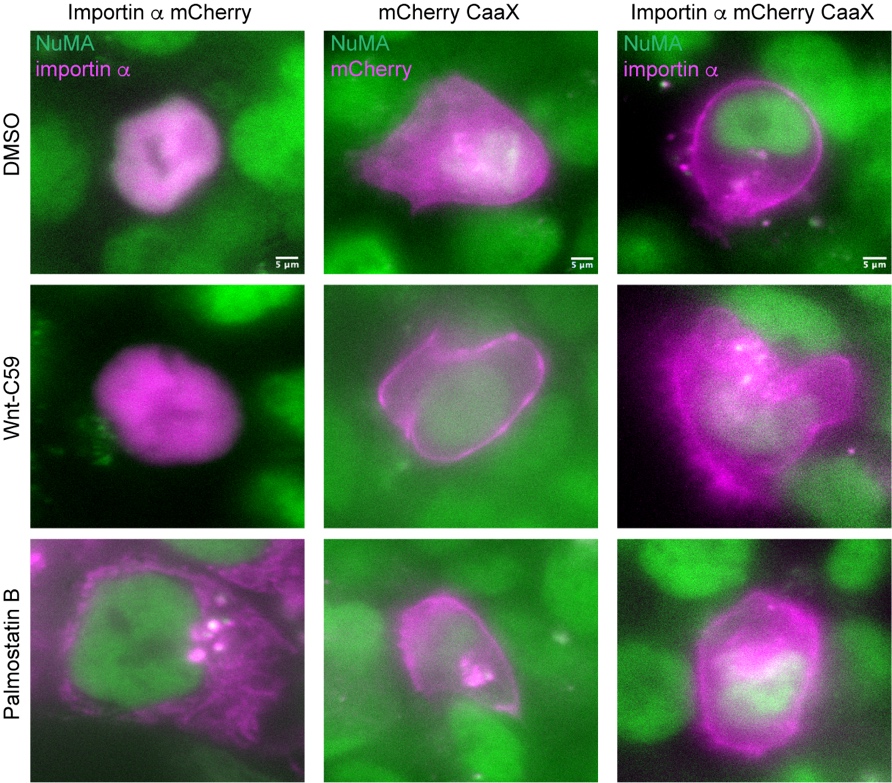

**Expanded View Figure 3. CaaX Modified Importin α Localizes to the Plasma Membrane Independent of Palmitoylation.**

Immunofluorescent images of HCT116 cells transfected with importin α-mCherry, mCherry-CaaX or importin α-mCherry-CaaX treated with DMSO, Wnt-C59 or palmostatin. Importin α-mCherry-CaaX localizes to the plasma membrane in all drug treatments. Scale bar=5μm.

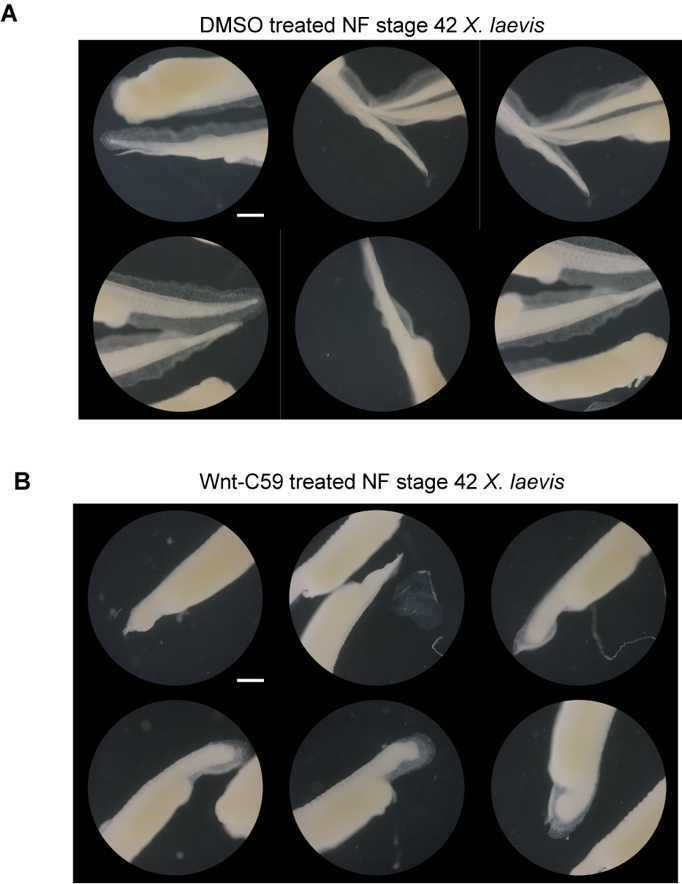

**Expanded View Figure 4. Wnt-C59 Treated *X. laevis* Tadpoles Exhibit Shortened Tail Length.**

A) Brightfield images of NF Stage 42 *X. laevis* tadpole tails treated with DMSO from 24hpf to 72hpf. DMSO treated tadpoles exhibit normal development and typical length tails. Scale bar=500μm.

B) Brightfield images of NF Stage 42 *X. laevis* tadpole tails treated with Wnt-C59 from 24hpf to 72hpf. Wnt-C59 treated tadpoles exhibit abnormal development, shortened tails, and abnormally shaped tails. Scale bar=500μm.

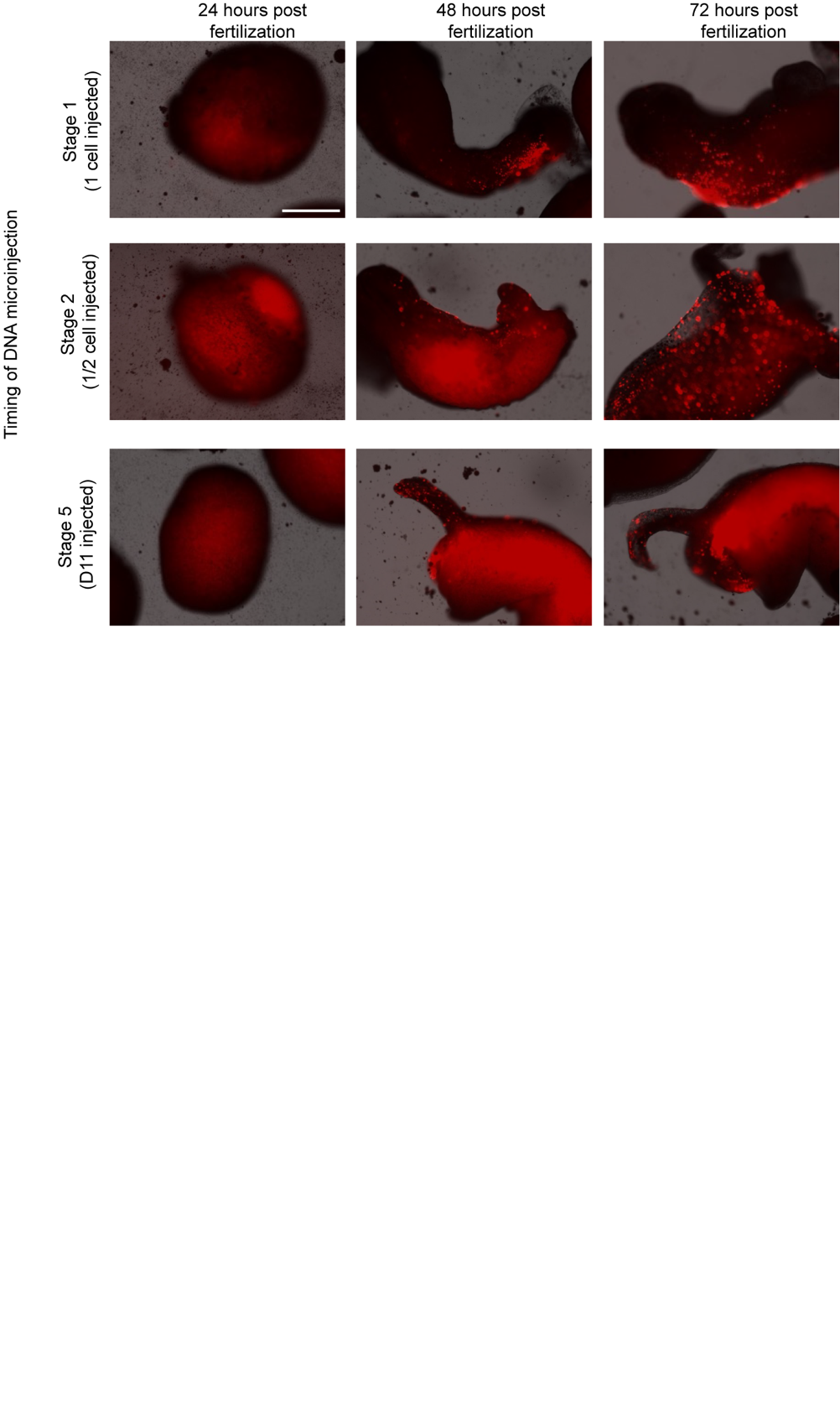

**Expanded View Figure 5. Importin α Overexpression Produces Severe Developmental Defects in *X. laevis* Embryos.**

Immunofluorescent images of *X. laevis* embryos co-injected with importin α-mCherry-CaaX pcDNA4TO and pcDNA6TR at 24, 48, and 72 hours post fertilization. Embryos were injected at either the 1 cell, 2 cell (injected into 1 of 2 cells), or 16 cell stage (injected into the D11 blastomere). All 1 and 2 cell injected embryos exhibit high mortality and severe developmental defects. D11 injected embryos had better survivability though some embryos still exhibited defects. Scale bar=500μm.
